# Atypical Neocortical Development in the *Cited2* Conditional Knockout Leads to Behavioral Deficits Associated with Neurodevelopmental Disorders

**DOI:** 10.1101/2020.09.02.279646

**Authors:** Nikolaus Wagner, Jessica L. MacDonald

**Affiliations:** Department of Biology, Program in Neuroscience, Syracuse University, Syracuse NY

**Keywords:** repetitive behaviors, ultrasonic vocalizations, sensory filtering deficits, somatosensory cortex, cortical dysfunction

## Abstract

The mammalian neocortex develops from a single layer of neuroepithelial cells to form a six-layer heterogeneous mosaic of differentiated neurons and glial cells. This process requires a complex choreography of temporally and spatially restricted transcription factors and epigenetic regulators. Even subtle disruptions in this regulation can alter the way the neocortex forms and functions, leading to a neurodevelopmental disorder. One epigenetic regulator that is essential for the precise development of the neocortex is CITED2 (<u>C</u>BP/p300 <u>I</u>nteracting <u>T</u>ransactivator with <u>ED</u>-rich termini). *Cited2* is highly expressed by intermediate progenitor cells in the subventricular zone during the generation of the superficial layers of the neocortex. A forebrain-specific conditional knockout of *Cited2* (cKO) exhibits reduced proliferation of intermediate progenitor cells embryonically, leading to reduced thickness of the superficial layers and a specific reduction in the somatosensory neocortical length postnatally. Further, the *Cited2* cKO displays decreased corpus callosum volume and dysregulation of precise neuronal connectivity within the somatosensory cortex. Here, we explore the behavioral consequences resulting from this aberrant neocortical development. We demonstrate that *Cited2* cKO mice display decreased maternal separation-induced ultrasonic vocalizations as neonates, and an increase in the repetitive behavior of rearing and lack of habituation following repeated acoustic startle as adults. They do not display alterations in anxiety-like behavior, overall locomotor activity or social interactions. Together with the morphological, molecular, and connectivity disruptions, these results identify the *Cited2* cKO neocortex as an ideal system to study mechanisms underlying neurodevelopmental and neuroanatomical disruptions with relevance to human neurodevelopmental disorders.

## Introduction

Pathophysiological disruptions in neuronal development and connectivity within the neocortex, the brain area responsible for high cognitive and associative functions, occur in multiple neurodevelopmental disorders. For example, a reduction in connectivity (Frazier and Hardan, 2009) and synchronization between the two cerebral hemispheres is common in autism spectrum disorders (ASD) (Dinstein et al., 2011). These interhemispheric connections are made by the axons of callosal projection neurons (CPN), a broad population of commissural neurons that are defined by the extension of an axon across the corpus callosum (CC), the largest axonal tract in the mammalian brain, to connect the two cerebral hemispheres. CPN are excitatory pyramidal projection neurons that are found throughout the neocortical layers (∼80% in layers II/III in mouse, 20% in V, and a small percent in VI), but that are the predominant subtype of projection neuron within layers II/III.

CPN are a heterogeneous population of interhemispheric neurons; they express unique combinations of genes that define them as a broad population, and other genes that identify distinct subpopulations within CPN during development (Molyneaux et al., 2009; Fame et al., 2011; MacDonald et al., 2013). For example, CPN within each functional neocortical area have distinct molecular and morphological characteristics (Lomber et al., 1994; Olivares et al., 2001; Grove and Fukuchi-Shimogori, 2003; Benavides-Piccione et al., 2006; O’Leary et al., 2007). Disruptions in CPN development and connectivity are associated with multiple neurodevelopmental disorders, including ASD (Egaas et al., 1995; Piven et al., 1997; Mukaetova-Ladinska et al., 2004; Herbert and Kenet, 2007; Frazier and Hardan, 2009; Srivastava et al., 2012), Rett syndrome (Belichenko et al., 1994; Armstrong et al., 1995; Kishi and Macklis, 2004, 2010; Wood et al., 2009; Ribeiro et al., 2020), ADHD (Hynd et al., 1991; Roessner et al., 2004; Seidman et al., 2005), and schizophrenia (Swayze et al., 1990; Tibbo et al., 1998; Innocenti et al., 2003; Wolf et al., 2008). CPN thus play key, diverse roles in complex associative and integrative cognition, and disruptions in their development profoundly alter behavior.

One critical regulator of neocortical CPN development and interhemispheric connectivity is the transcriptional co-regulator CITED2 (Fame et al., 2016). CITED2 acts as a transcriptional co-activator by recruiting the CBP/p300 histone acetyltransferase complex to activate gene transcription (Bhattacharya et al., 1999; Yahata et al., 2000; Bragança et al., 2003; Chou et al., 2012), or as a transcriptional co-repressor by preventing transcription factors from binding to CBP/p300 (Bhattacharya et al., 1999; Yoon et al., 2011). *Cited2* is expressed broadly by progenitors of the embryonic (E)15.5 subventricular zone, during the peak of superficial layer CPN birth, with a progressive post-mitotic refinement in expression to CPN of somatosensory cortex postnatally (Fame et al., 2016). There is a broad reduction of TBR2-positive progenitors at E15.5 across the neocortex in the *Cited2* forebrain-specific conditional knockout (cKO; Emx1-mediated), resulting postnatally in both reduced thickness of superficial layers and a highly area-specific reduction of layer II/III somatosensory neocortical length. Further, the *Cited2* cKO displays reduced interhemispheric connectivity throughout the neocortex, and disorganized axonal and dendritic connectivity specifically within somatosensory cortex. Strikingly, a NEX-Cre post-mitotic forebrain-specific *Cited2* cKO identified that *Cited2* is not required post-mitotically for these phenotypes, even though some processes, like arealization and dendritic arborization, are completed post-mitotically (Fame et al., 2016). This indicates that CITED2 functions within neocortical progenitors to both broadly regulate generation of superficial layer CPN throughout the neocortex, and in an areally-restricted manner to refine the distinct identity and precise connectivity of somatosensory CPN.

Deficits in proliferation and differentiation of neural progenitor cells are a major convergence point for a range of neurodevelopmental disorders, including ASD and intellectual disability (Ernst, 2016), and the *Cited2* cKO neocortex displays disruptions in both of these processes (Fame et al., 2016). Thus, we asked the question of whether these developmental disruptions result in behavioral alterations that recapitulate aspects of human neurodevelopmental disorders. We used a multitude of behavioral assays to measure verbal communication, locomotor activity, restrictive and repetitive behaviors, anxiety-like behavior, social interactions, and sensorimotor gaiting to compare *Cited2* cKO mice (Emx1-mediated) with their wild-type (WT) and heterozygous (HET) littermates. We found that *Cited2* cKO mice display deficits in maternal separation-induced ultrasonic vocalizations as neonates, and increased restrictive and repetitive behaviors and sensitization to acoustic startle as adults. They do not, however, display any observable differences in locomotor activity or social interactions. Thus, the aberrant neocortical development identified in the *Cited2* cKO leads to life-long behavioral disruptions associated with neurodevelopmental disorders, identifying the *Cited2* cKO neocortex as a novel system to study mechanisms underpinning atypical neurodevelopment and neurodevelopmental disorders.

### Experimental Procedures

#### Mice

All animal experimental protocols were approved by the Syracuse University Institutional Animal Care and Use Committee and adhere to NIH ARRIVE guidelines. Mice were group housed at a maximum of 5 mice per cage on a 12:12 h light/dark cycle, and were given food and water *ad libitum. Cited2* conditional floxed mice (C2f) (Preis et al., 2006) were generously provided by Dr. Sally Dunwoodie, Victor Chang Cardiac Research Institute, University of New South Wales, Australia. They were rederived by The Jackson Laboratory (Bar Harbor, Maine, USA) using JR 664 C57BL/6 oocytes to generate live mice at an SPOF health status. Emx1-Cre mice (Guo et al., 2000) were obtained from The Jackson Laboratory (RRID:IMSR_JAX:005628). To avoid non-specific cre recombinase activity in oocytes (Hayashi et al., 2003), all conditional knockouts were generated by crossing C2f fl/fl females with C2f fl/+; Emx1 cre+ males, and no offspring from C2f fl; Emx1 cre+ dams were analyzed. The morning of the day of the appearance of the vaginal plug was defined as E0.5. The day of birth was designated P0. Genotypes were assessed by PCR on genomic DNA extracted with the KAPA extract kit using the following primers: *Cited2* flox/flox, flox/wt, and wt/wt were determined using – *Cited2* Forward 5’-GTC TCA GCG TCT GCT CGT TT-3’; *Cited2* Reverse 5’-CTG CTG CTG TTG GTG ATG AT -3’. Emx1 was distinguished from Emx1-Cre using – Emx1 WT Forward 5’-GAA GGG TTC CCA CCA TAT CAA CC-3’; Emx1 WT Reverse 5’-CAT AGG GAA GGG GGA CAT GAG AG-3’; Emx1-Cre Reverse 5’-TGC GAA CCT CAT CAC TCG TTG C-3’.

#### Immunohistochemistry

Immunohistochemistry was performed as previously described (Fame et al., 2016). Briefly, brains were post-fixed overnight in 4% PFA/PBS at 4°C, then were sectioned on a VT1000S vibrating microtome (Leica Microsystems). Sections were incubated in primary antibody dilutions at 4°C overnight, and appropriate secondary antibodies were selected from the Molecular Probes Alexa series (Invitrogen, Carlsbad, CA). Antigen retrieval methods were required to expose antigens for some of the primary antibodies. Sections were incubated in 0.1M citric acid (pH=6.0) for 10 min. at 95-98°C. Primary antibodies were used as follows: rat anti-CTIP2 (Abcam Cat# ab18465, RRID:AB_2064130), rabbit anti-β-galactosidase (Thermo Fisher Scientific Cat# A-11132, RRID:AB_221539), mouse anti-SATB2 (Abcam Cat# ab51502, RRID:AB_882455), rabbit anti-TBR1 (Abcam Cat# ab31940, RRID:AB_2200219), rabbit anti-BHLHB5 (Abcam ab204791), mouse anti-NeuN (Millipore Cat# MAB377, RRID:AB_2298772). Secondary antibodies from Invitrogen Alexa Series (Invitrogen).

#### Quantification of Neocortical Length and Thickness

All length and thickness measurements were performed with images of matched sagittal 50μm sections using ImageJ to trace the curvature of the neocortical surface. The somatosensory area was identified by the expression of Bhlhb5 in layers II/III. Deep layers (V-VI) were identified as those including cells expressing high levels of CTIP2 and deeper. Superficial layers (II-IV) were identified as those superficial to high level CTIP2 expression.

#### Behavior

For all behavioral tests, mice were allowed to acclimate to the testing room for a 30-minute period. Each test was run at the same time of the day for all litters by the same experimenter blind to genotype. Only one assay was preformed per day starting when mice reached 10 weeks of age. More specific methods for each test are as follows:

### Ultrasonic Vocalizations

After acclimation, pups were isolated from their dam into a prewarmed cage. Pups were then individually placed into a sound attenuation chamber and their USVs were recorded with an ultrasonic microphone (Patterson M500-384) over a period of five minutes. After this five-minute period, the neonates were returned to their prewarmed cage. Number of calls, average duration of calls, and peak frequency of calls were measured using Noldus UltraVox XT software. Individual USVs were counted if they met the following criteria; frequency was between 30kHZ and 125kHz, duration was between 5ms and 500ms with a gap of ≥6ms, and amplitude was greater than 250a.u. above background. It is important to note there is no observable difference in the size of pups based on genotype. As this is a forebrain specific cKO, neonates are expected to have no difference in lung capacity or physical ability to make calls.

### Open Field Maze

An open-field maze using a photo-beam recording system (SD Instrument) was used to record activity in a novel clear box (40 cm × 40 cm) for 30 minutes with a background white noise of ∼70 dB and light intensity level of ∼1,000 Lux. The mouse’s travel distance, travel speed, time spent resting, and rearing were recorded and quantified using SanDiego’s PAS Reporter software.

### Light-Dark Box

Mice were placed facing the same corner of a box consisting of a light side measuring (40cm x 26 cm) with an illumination level of ∼1,000 lux and a dark side measuring (40cm x 18cm) with an illumination level of <20 lux. Mice were removed after 10-minutes and distance traveled in the light portion, time in each side of the box, and number of transitions were recorded using the Ethovision XT software (Noldus).

### Elevated-Plus Maze

A plus shaped maze elevated 63.5cm off the ground with two open arms (35cm x 5cm) and two closed arms of the same size but enclosed by walls 20cm high with no ceiling was employed. Mice were placed in the center of the maze facing one of the open arms; their activity was recorded over a 5-minute period. Using Ethovision XT (Noldus), we measured the number of entries into each arm, time spent in each arm, distance traveled, and amount of time with their nose reaching over the edge of the open arms.

### Social Preference Test

Social interactions were evaluated using the three-chambered social approach test as described (Yang et al., 2011). The apparatus consists of three chambers (each measuring 40cm ⨯20cm) separated by walls with removable doors. After a 5-minute acclimation period in the center chamber with the doors closed, the doors were opened, and the mouse was allowed to explore all three empty chambers for ten minutes. The mouse was then placed back in the closed center chamber and two identical wire cups were placed in the left and right side, one containing an age and sex matched novel mouse and the other left empty as a novel object. The doors were opened again, and the mouse was free to explore for another 10 minutes. For this third portion, we used Ethovision XT (Noldus) to assess total distance traveled, amount of time in each chamber, number of entries to each chamber, and time sniffing within 2cm of the wire cup.

### Startle Response, Pre-pulse Inhibition, and Habituation

To measure sensorimotor gating and sensory filter we employed a three-block assay (SR-Lab; SD instruments) as described previously (Conti, 2012). Prior to the start of the tests, there was a 5-minute acclimation period in the chamber with a 70dB background noise. The first block of the test, used to further acclimate and assess individual startle response (SR), consisted of six trials of a 120dB startling stimulus presented for 40ms. All trials contained a randomized 10-30s gap between stimuli that averaged to 20s. The second block was used to calculate pre-pulse inhibition (PPI) and consisted of each of the following stimuli presented in a pseudorandomized order; twelve stimulus alone (120dB) trials, twelve trials with a pre-pulse preceding the 120dB startle stimulus by 100ms (6, 12, 15, or 18dB above background), and eight trials with no stimulus to assess for random baseline activity. Block three was a repeat of block 1 (six, 120dB alone startle stimuli) to assess for habituation and sensitization. SR for each mouse was calculated by averaging the SR for each of the six trials in Block 1. Percent prepulse inhibition was calculated as 100-100*(average startle amplitude with the prepulse/average startle amplitude of the Block#2 startle stimulus alone trials). Percent habituation/sensitization was calculated as 100*((average startle amplitude of Block #1-average startle amplitude of Block #3)/average startle amplitude of Block #1).

### Statistical Analyses

GraphPad Prism 8.0 (GraphPad Software) was used to carry out the statistical analyses. No statistical methods were used to predetermine sample sizes, but our sample sizes are similar to those generally employed in the field. Our statistical tests consisted of one-way ANOVA with Tukey’s multiple comparison to determine statistical significance between groups. Sample size and statistical test are specified in each figure legends.

## Results

### Cited2 regulates neocortical size, including superficial layer thickness and somatosensory length

CITED2 is a key regulator of development of many organs, including eye, liver, kidney, and heart, and processes including neural tube closure; thus, the *Cited2*-null is embryonic lethal, exhibiting multiple developmental defects (Barbera et al., 2002; Weninger, 2005; Qu et al., 2007; Val et al., 2007; Chen et al., 2008; Xu et al., 2008). To bypass these early developmental disruptions, including exencephaly, we generated forebrain-specific *Cited2* cKO mice using Emx1-promoter driven cre-recombinase (Emx1-Cre; (Guo et al., 2000)) and the *Cited2 floxed* line (Preis et al., 2006). We first determined that the resulting offspring have the same phenotype as previously published (Fame et al., 2016). Successful excision of the *Cited2* gene was verified via expression of a β-galactosidase reporter, identified by immunohistochemistry; *Cited2* cKOs display high β-galactosidase expression restricted to the forebrain while no β-galactosidase is detected in wildtype littermates (Fig. 1*A*). We employed immunohistochemistry and verified the reduced thickness of the superficial layers of the somatosensory cortex, identified as SATB2 positive cells basal to the layer V marker CTIP2 (Fig. 1*B-D*). We additionally replicated the previous finding of a significant reduction in the length of the somatosensory cortex using immunohistochemistry for Bhlhb5 to identify and measure the length of layer II/III somatosensory cortex, on sagittal sections carefully matched by anatomical landmarks (Fig. 1*E-F*). We were also able to validate the diminished corpus callosum (CC) size in *Cited2* cKOs by evaluating the cross-sectional area of the CC (Fig. 1*G*). Together, these data confirm that our *Cited2* cKO mice display morphological phenotypes that are consistent with previous findings, and mirror morphological deficits observed in a variety of neurodevelopmental disorders.

**Figure 1.**
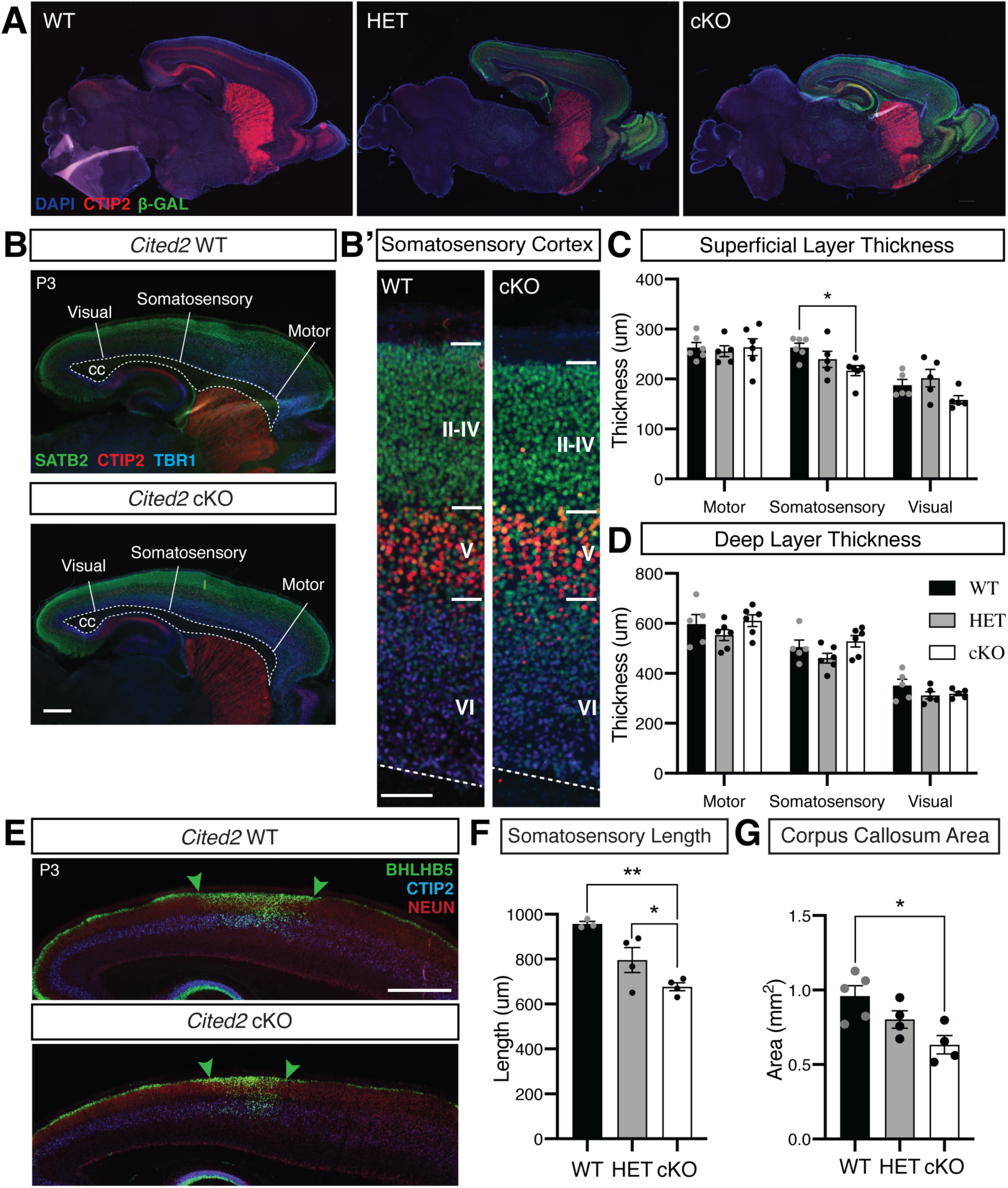
Conditional loss of *Cited2* leads to disrupted neocortical development and morphological abnormalities associated with neurodevelopmental disorders. ***(A)***, *Cited2* excision, driven by an Emx1 promoter, is visualized by expression of a β-GAL reporter (green). β-gal expression is restricted to the forebrain of *Cited2* cKO and HET mice at postnatal day (P) 3. ***(B-D)***, Neocortical laminar thickness was measured in the visual, somatosensory, and motor areas of the P3 neocortex. Superficial layers (II-IV) were identified by the expression of SATB2 (green) in neurons located basally to the expression of CTIP2 (red) in layer V. TBR1 (blue) identifies layer VI. *Cited2* cKO mice display decreased laminar thickness of the somatosensory cortex (***B’***), generated by a decrease in thickness of the superficial layers (II-IV) with no change in deep layers thickness (V-VI) (n = 5-6 mice per genotype). **(*E-F)***, Loss of *Cited2* leads to decreased somatosensory areal length, as identified by the expression of BHLHB5 (green) in the superficial layers. Green arrows mark the boarders of the somatosensory cortex. ***G***, *Cited2* cKO mice have a decreased corpus callosum (CC) cross-sectional area compared to their WT littermates. The CC is labeled in both images in ***B***. \**p* < 0.05; \*\**p* < 0.01 (One-way ANOVA with Tukey’s post hoc analysis). Error bars denote s.e.m. Scale bars: ***A*** and ***E***, 1000μm; ***B***, 500μm; ***B’***, 100μm.

### Cited2 conditional knockout leads to disruptions in neonatal ultrasonic vocalizations

Having verified that *Cited2* cKOs display specific disruptions in neocortical development, we performed a behavioral characterization for traits of neurodevelopmental disorders associated with these morphological disruptions. Deficits in verbal communication is a common phenotype of many neurodevelopmental disorders. Neonatal mouse verbal communication can be induced and measured using a maternal separation paradigm where pups emit retrieval calls in ultrasonic frequencies (Scattoni et al., 2009). Deficits in early postnatal maternal separation-induced ultrasonic vocalizations (USVs) have been identified in multiple murine models of neurodevelopmental disorders, including the *Shank, Pten*, and *Kirrel3* models of ASD (Wöhr et al., 2011; Binder and Lugo, 2017; Hisaoka et al., 2018), the *CBP* model of Rubinstein-Taybi Syndrome (Wang et al., 2010), and the methylazoxymethanol acetate induced model of schizophrenia (Potasiewicz et al., 2019), among others. We used a maternal separation assay to evoke USVs in our *Cited2* cKO mice. Ultrasonic spectrograms allow us to visualize evoked calls, and show a clear decrease in the number of calls made by *Cited2* cKO mice (Fig. 2*A*). When these calls are quantified over 5 minutes of testing, male *Cited2* cKO mice display decreased total number of USVs at P5 and P7 compared to WT, while female *Cited2* cKOs show a significant reduction at P5 and a trend to reduced calls at P7 (p=0.069; unpaired t-test) (Fig. 2*B*). Although not significantly different from either genotype, it is interesting that HET mice appear to have an intermediate phenotype. Neither sex nor genotype have an effect on call duration (Fig. 2*C*) or average peak frequency (data not shown), suggesting *Cited2* cKO and HET call numbers are not a result of physical limitations.

**Figure 2.**
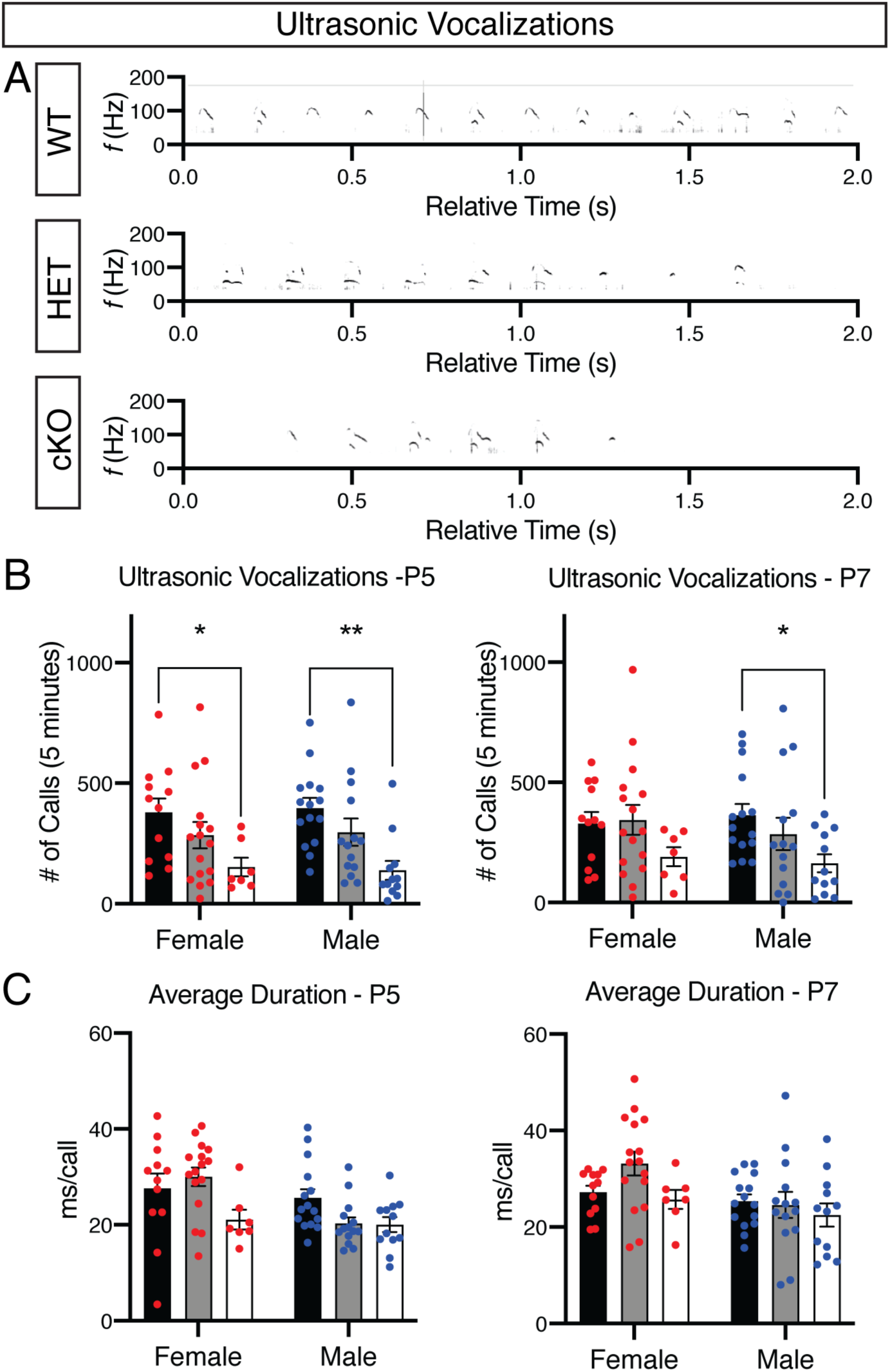
Forebrain-specific loss of *Cited2* leads to deficits in maternal separation-induced ultrasonic vocalizations in neonates. (***A)***, Representative spectrograms of segments from 5-minute recordings of ultrasonic vocalization (USV) from *Cited2* WT, HET, and cKO P5 neonatal mice following maternal separation. (***B)***, Quantification of the number of USVs over the full 5-minute period after maternal separation shows a significantly decreased number of calls for *Cited2* cKO males and females compared to WT littermates at P5. At P7, *Cited2* cKO males display significantly decreased number of calls while females show a trend towards decreased calls. Despite having a decreased number of calls, no differences were detected in average duration per call ***(C***). N = 7-15 per sex and genotype. Females are represented by red dots and males by blue dots. \**p* < 0.05; \*\**p* < 0.01 (One-way ANOVA with Tukey’s post hoc analysis)

### Cited2 conditional knockout leads to increased repetitive behaviors, without a change in overall locomotor behavior

Restricted and repetitive behaviors are another persistent phenotype characterized in ASD and individuals with other neurodevelopmental disorders. These behaviors manifest themselves in murine models of neurodevelopmental disorders in multiple ways, including alterations in rearing activity (Kim et al., 2016), which have been reported in *Shank, Pten*, and *CBP* neurodevelopmental disorder models (Lugo et al., 2014; Lee et al., 2015; Zheng et al., 2016). To evaluate rearing in *Cited2* cKO model, we measured vertical beam break activity during a 30-minute open-field test (San Diego Instruments). Both female and male *Cited2* cKOs have significantly increased rearing compared to HET and WT (Fig. 3*A*). Increased rearing may be viewed as a result of hyperactivity; however, *Cited2* cKOs show no locomotor differences in the open-field maze measured through total distance traveled, average speed while moving, average resting time, and entries to the center of the maze (Fig. 3*B-E*). Taken together, increased rearing patterns of *Cited2* cKO mice mirror the restricted and repetitive behaviors phenotype demonstrated by other neurodevelopmental disorder murine models.

**Figure 3.**
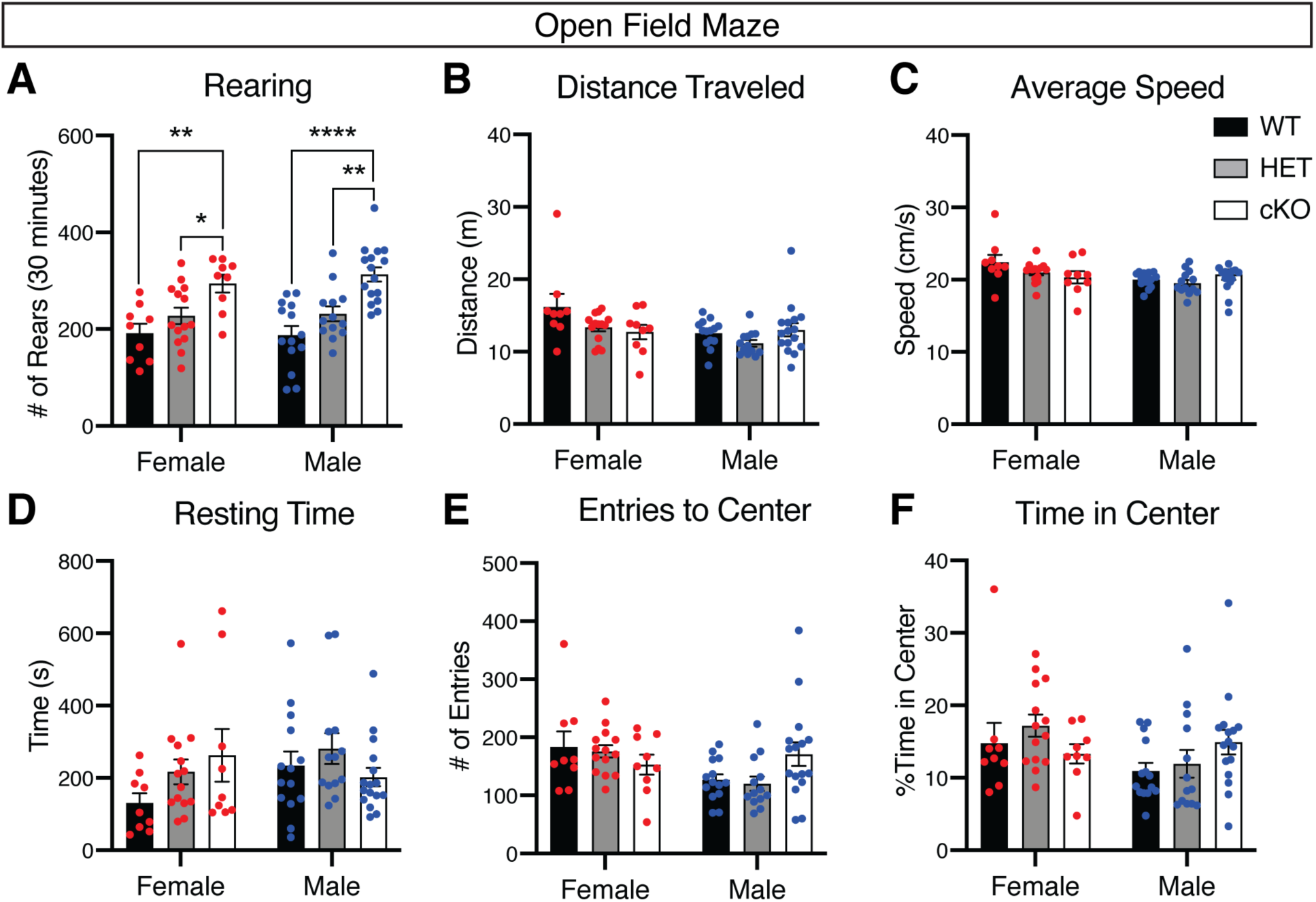
*Cited2* cKO mice display increased repetitive behaviors with no change in locomotor activity in the open-field maze. (***A)***, During the 30 minute Open-Field Test, *Cited2* cKO mice of both sexes reared significantly more than WT mice, measured by vertical beam breaks. Despite this increase in vertical activity, *Cited2* cKO mice do not display a hyperactive phenotype or altered locomotor activity as evidenced by no observed differences in total distance traveled ***(B)***, average speed while moving ***(C)***, resting time ***(D)***, or number of transitions between the outside and center of the maze ***(E). F***, *Cited2* cKO mice do no spend more or less time in the center of the maze, suggesting no alterations in anxiety-like phenotypes. N = 9-16 mice per sex and genotype. Females and males are represented by red and blue dots, respectively. \*\**p* < 0.01; \*\*\*\**p* < 0.0001 (One-way ANOVA with Tukey’s post hoc analysis)

### Cited2 conditional knockout does not lead to a significant change in anxiety or social behaviors

Anxiety related disorders have an approximate 40% comorbidity rate with neurodevelopmental disorders such as ASD and Schizophrenia (Braga et al., 2013; Zaboski and Storch, 2018). Anxiety-like behavior in mice is evaluated using a variety of assays including the open-field maze, elevated plus maze, and the light-dark box (Lezak et al., 2017). We utilized these three assays to further characterize the behavior of *Cited2* cKO mice and assess for anxiety-like behaviors. We observe no difference in anxiety-like behaviors of *Cited2* HET or cKO mice in the open field maze, measured by the amount of time the spent in the center of the maze (Fig. 3*F*). For the elevated plus maze and the light-dark box, both *Cited2* cKOs and HETs behave like WT, showing no anxiety-like phenotypes in either assay (Fig. 4*A-B*). Of note, we observe no differences in the number of transitions between the light and dark side (Fig. 4*B*), as well as no differences in total distance traveled for either test (data not shown), again indicating that the mice do not display hyperactivity.

**Figure 4.**
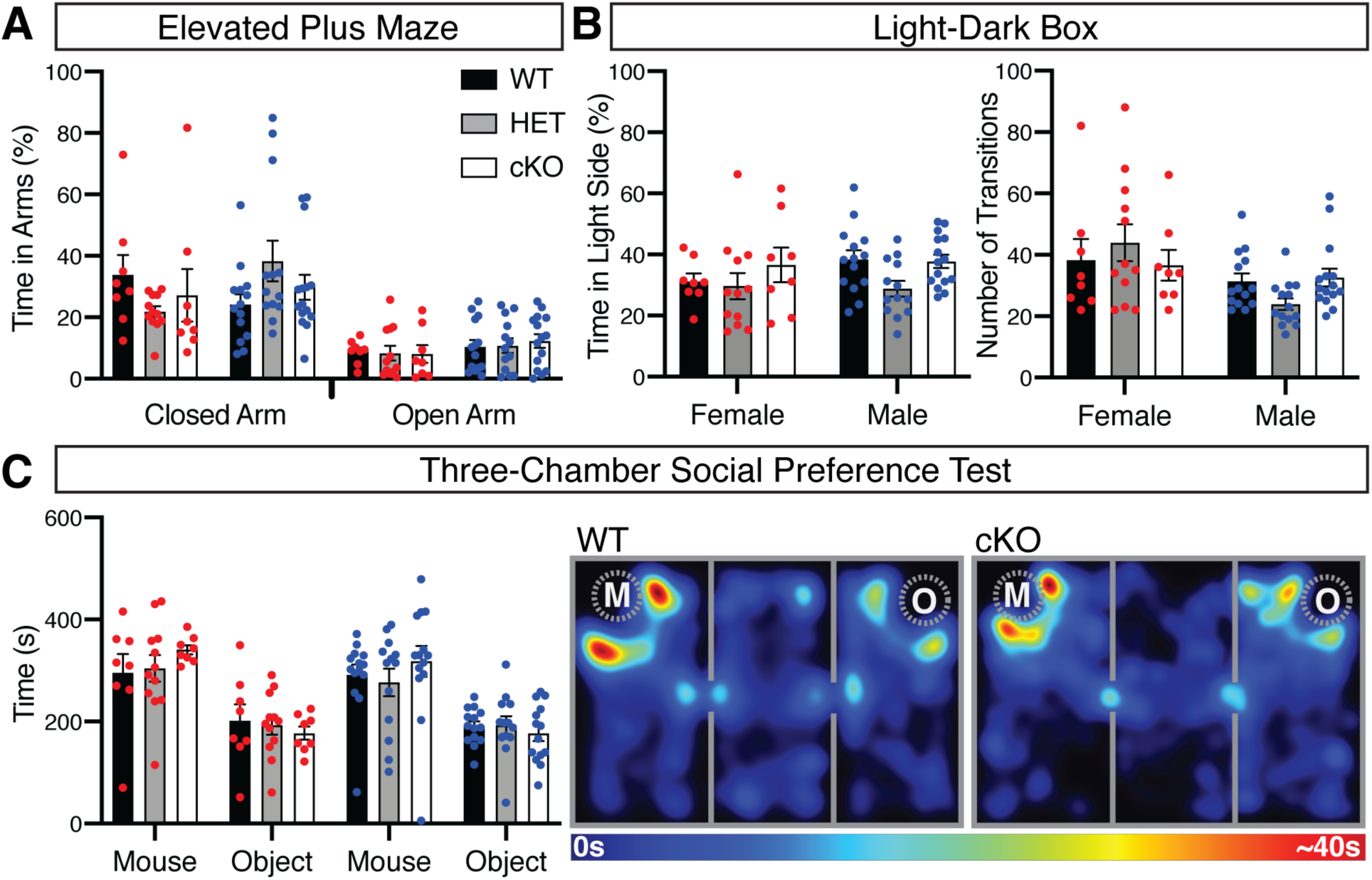
*Cited2* cKO mice do not display alterations in anxiety-like behaviors or social interactions. *Cited2* cKO animals perform comparable to WT in assays that measure anxiety-like behaviors including the classic Elevated-Plus maze and the Light-Dark Box. (***A)***, During the 5-minute Elevated Plus test, *Cited2* cKO mice behaved similar to WT in that they spent more time in the closed arm than the open. (***B)***, *Cited2* WT, HET, and cKO mice all spent similar time in the light portion of the Light-Dark box indicating no alterations in anxiety-like behavior. All three genotypes displayed a comparable number of transitions between the light and dark portions of the box, indicating similar overall activity levels. (***C)***, *Cited2* cKO mice demonstrated similar sociability to WT in the Three-Chamber Social Preference Test, with all three genotypes spending more time with a novel mouse, quantified in the bar graph and visualized with the representative heatmap of WT and *Cited2* cKO males. M, novel mouse; O, Novel Object. N = 8-14 mice per genotype and sex. Females and males are represented by red and blue dots, respectively. One-way ANOVA with Tukey’s post-test was used to determine statistical significance.

A third common phenotype of many neurodevelopmental disorders is a deficit in social interactions. In mouse models, social interactions are often evaluated using the three-chamber social preference test (Yang et al., 2011). We used that assay to explore potential differences in social interactions due to *Cited2* loss-of-function. Interestingly, *Cited2* cKO and HET mice perform similarly to WT, spending more time with the novel mouse than the novel object, indicating no alteration in social preference (Fig. 4*C*).

### Cited2 conditional knockout causes a significant disruption in auditory startle habituation]

Prepulse inhibition (PPI) is a measure of sensorimotor gating, which is disrupted in neurodevelopmental disorders such as schizophrenia and some cases of ASD. Individuals diagnosed with schizophrenia typically show decreased PPI and thus impaired sensorimotor gating (Braff et al., 2001), a phenotype that is mirrored in murine models of schizophrenia (Powell et al., 2009). The results of startle response and PPI tests on patients with other neurodevelopmental disorders are less clear. For example, it has been reported that ASD individuals may not have altered PPI, but have increased sensitization and altered habituation (Madsen et al., 2014; Sinclair et al., 2017). To assess sensorimotor gating, startle response, and habituation in *Cited2* cKO mice, we used an acoustic startle paradigm consisting of three blocks. Block#1 and Block#3 contain 120dB stimuli repeated six times to evaluate startle response and habituation, while Block#2 contains six different stimuli presented in a pseudo randomized order to evaluate pre-pulse inhibition (Fig. 5*A*). *Cited2* cKO mice show no significant deficits in PPI when compared to their WT littermate controls (Fig. 5*B-C*). Although the *Cited2* cKOs do not show a significant difference in initial startle response to the auditory stimuli (Fig. 5*D*), *Cited2* cKOs of both sexes show a significantly decreased habituation, indicating an enhanced sensitization to auditory stimuli throughout the testing and deficits in sensory filtering (Fig. 5*E*). Interestingly, the HETs again appear to have an intermediate phenotype, showing neither the habituation observed in the WTs nor the sensitization observed in the *Cited2* cKOs. Taken together, *Cited2* cKO mice display deficits in sensory filtering evident through decreased habituation / enhanced sensitization, but no observable difference in sensorimotor gating measured through PPI.

**Figure 5.**
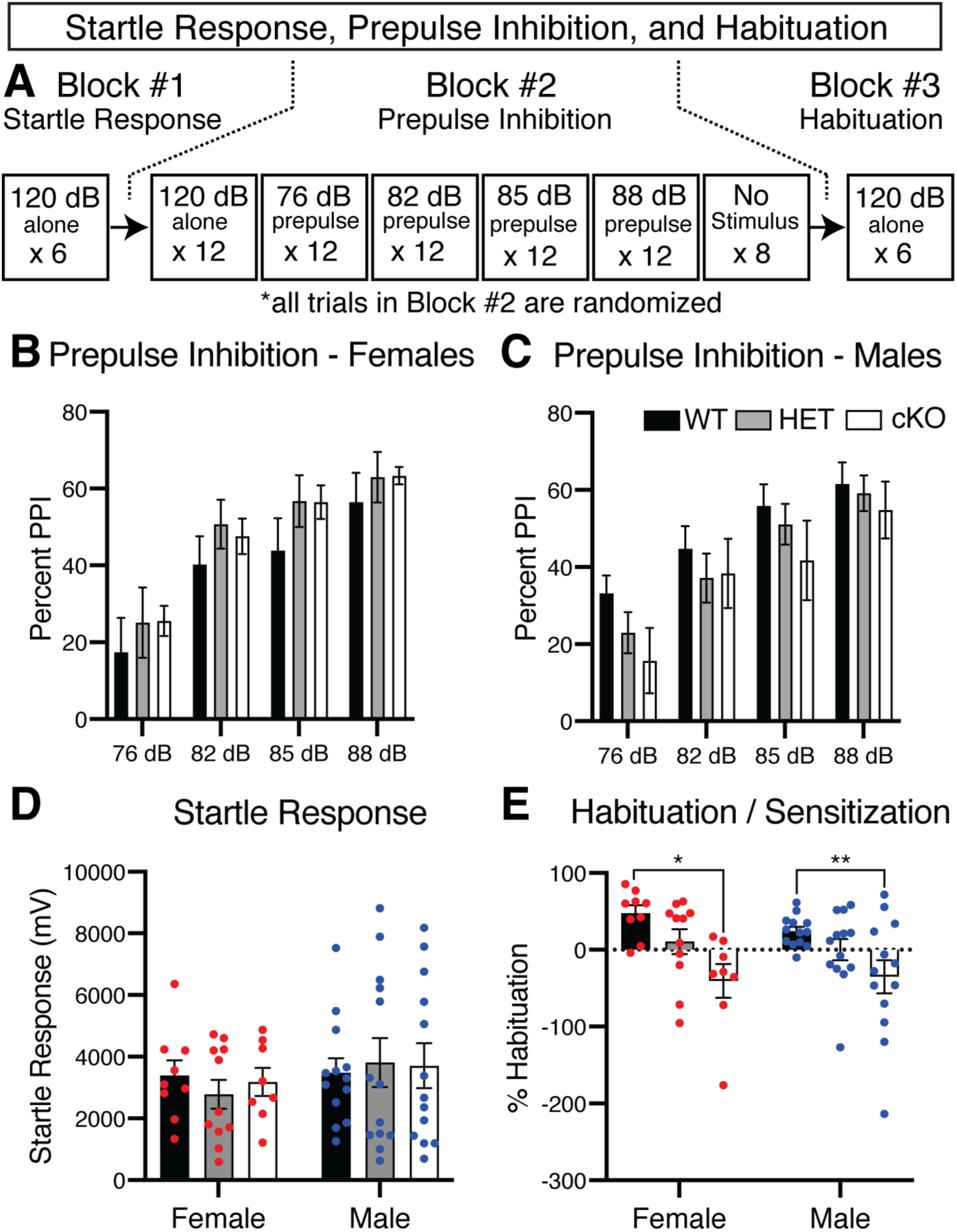
*Cited2* cKO mice display decreased habituation with no change to startle response or pre-pulse inhibition. (***A)***, To assess startle response, pre-pulse inhibition, and habituation/sensitization, we employed a three-block assay using the San Diego SR lab apparatus. The first and third block consisted of a 120dB acoustic tone repeated 12 times to measure startle response and habituation/sensitization, the second consisted of six different types of stimuli presented in a pseudo random order to assess pre-pulse inhibition. (***B)***, Female *Cited2* HET and cKO mice displayed no differences in pre-pulse inhibition (PPI) compared to WTs while males ***(C)*** display trends towards decreased PPI with lower pre-pulse levels; however, the trends are not statistically significantly. (***D)***, We observed no genotype or sex difference for average startle response, however, (***E***) *Cited2* cKO mice of both sexes demonstrated decreased habituation across the entire test. On average, their response to acoustic stimulus in Block #3 was greater than their response in Block #1, suggesting they became more sensitive to the acoustic startle as the test progressed. N = 8-15 per sex and genotype. Females and males are represented by red and blue dots, respectively. \**p* < 0.05; \*\**p* < 0.01 (One-way ANOVA with Tukey’s post hoc analysis)

## Discussion

Taken together, our data demonstrate that the atypical neuronal development in the *Cited2* cKO neocortex is sufficient to cause behavioral deficits that mirror phenotypes associated with many neurodevelopmental disorders. Chief among these deficits are alteration in verbal communication, seen through neonatal maternal separation-induced ultrasonic vocalizations; increased restrictive and repetitive behaviors, measured through repetitive rearing; and decreased habituation to repetitive acoustic stimuli, suggesting inhibited sensory filtering. Although *Cited2* cKO mice display increased rearing, their overall activity is not affected. Locomotor activity, social interactions, anxiety like behaviors, and pre-pulse inhibition are also unaffected, indicating that forebrain specific *Cited2* loss-of-function disrupts some behaviors while leaving others unaltered. It is interesting to note that the *Cited2* HET mice demonstrate an intermediate phenotype between WT and cKO for most morphological and behavioral assays. Heterozygous human *CITED2* mutations are linked to congenital heart disease, suggesting haploinsufficiency (MacDonald et al., 2008). Our data suggest that *CITED2* haploinsufficiency could also disrupt neocortical development and behavior.

The Emx1-mediated *Cited2* cKO specifically disrupts *Cited2* expression in projection neurons and astrocytes of the forebrain, leaving expression intact in other cellular populations. Altered mouse behaviors have been shown to be affected by manipulating one particular subtype of neuron. For example, *Pten* mutant mice display deficits in social interactions, locomotor activity, and anxiety-like behavior, which can be improved with the transplantation of wild-type interneurons (Southwell et al., 2020). Although *Cited2* is highly expressed in interneuron progenitors (Fame et al., 2016), the Emx1-Cre conditional knockout does not disrupt its expression in this population, which may explain why we don’t see disruptions in these specific behaviors. Many mouse models of neurodevelopmental disorders have mutations in specific neuronal subtypes that differentially affect behaviors depending on what brain region is altered. For example, loss-of-function of *Cul3* in the prefrontal cortex corresponds with social behavioral deficits, while loss-of-function in the striatum contributes to repetitive behaviors (Rapanelli et al., 2019). The specific behavioral disruptions found in the *Cited2* cKO could provide additional information on the role of distinct neuronal subtypes or circuits in behaviors associated with neurodevelopmental disorders.

CITED2 acts as a transcriptional co-activator by recruiting the histone acetyltransferase (HAT) complex CBP/p300 (Bhattacharya et al., 1999). In humans, mutations in *CBP* or *p300* lead to Rubinstein-Taybi syndrome, a neurodevelopmental disorder characterized by moderate to severe intellectual disability, and facial and skeletal abnormalities (McArthur, 1967; Fred Petrij et al., 1995). *CBP* and *p300* mutant mice have overlapping phenotypes with the *Cited2* cKO, including decreased progenitor proliferation, decreased USVs, and increased rearing (Wang et al., 2010; Zheng et al., 2016). Thus, *Cited2* cKO might recapitulate a subset of this disorder, reflecting the restricted expression of *Cited2* to neocortical intermediate progenitors and CPN in comparison to the broad expression of *CBP* and *p300* in the brain.

We verified that the *Cited2* cKO has a reduced corpus callosum, consistent with the overall reduction in superficial layer thickness and CPN number. Further, *Cited2* cKO somatosensory CPN have been shown to project contralateral axons imprecisely, with a bimodal distribution of axonal projections covering a more expansive target area on the contralateral hemisphere than the tightly delineated homotopic areas targeted by wildtype CPN. HARDI diffusion spectral imaging (DSI) tractography analysis of interhemispheric connections also appears to show a significant reduction in projections connecting the somatosensory and premotor cortex (Fame et al., 2016). Deficits in connectivity and functionality of ipsilaterally connected brain regions have been well established in a variety of neurodevelopmental disorders including ASD, ADHD, and Schizophrenia. Of these regions, the prefrontal cortex (PFC) is most frequently reported as having deceased functional connectivity with other brain regions in ASD (Gotts et al., 2012; Rudie et al., 2012; Nair et al., 2013; Starck et al., 2013; Rane et al., 2015). Decreased connectivity with the PFC is further correlated with restrictive and repetitive behaviors, a keystone phenotype in the diagnosis of ASD (Delbruck et al., 2019). PFC functionality deficits are prevalent in schizophrenia as well, with both hypoactivation phenotypes (Huang et al., 2010; Denardo et al., 2015) and consistent reduced connectivity reported (Woodward et al., 2012; Orban et al., 2017; Giraldo-Chica et al., 2018; Chen et al., 2019; Bulletin et al., 2020). Although additional experiments are necessary to investigate intrahemispheric connectivity disruptions in the *Cited2* cKO, including those to the PFC, it is likely that disruptions in both long-distance intrahemispheric and interhemispheric connectivity contribute to the behavioral disruptions in the *Cited2* cKO.

It is interesting to note that in global *Cited2*-null mice, maternal folic acid supplementation prevents neural tube closure defects in offspring (Barbera et al., 2002), while maternal high-fat diet doubles the penetrance of heart defects and induces cleft palate in offspring (Bentham et al., 2010). Importantly, maternal folic acid supplementation in humans is associated with reduced risk of a neurodevelopmental disorder in offspring (Surén et al., 2013; Gao et al., 2016; Irwin et al., 2016). In contrast, human maternal obesity and/or high gestational weight gain is associated with higher probability of a neurodevelopmental disorder in offspring (Krakowiak et al., 2012; Rivera et al., 2015; Connolly et al., 2016). Folate receptor antibodies, which can block folate transport and lead to folate deficiency, are also prevalent in human ASD (Frye et al., 2013), and can induce communication, learning, and cognitive deficits in rodent models when administered during gestation (Desai et al., 2017). Further, maternal high-fat diet induces social behavior deficits in rodent models (Kang et al., 2014; Buffington et al., 2016). Whether or not maternal folic acid supplementation and high-fat diet can reduce or exacerbate, respectively, the *Cited2* cKO cortical phenotypes and aberrant behaviors is not known. However, the *Cited2* cKO could provide a novel system in which to test how *in utero* environmental exposures could lead to altered neocortical development and life-long alterations in behavior that recapitulate aspects of human neurodevelopmental phenotypes.

## Acknowledgements

The *Cited2*flox transgenic mice were generously provided by Dr. Sally L. Dunwoodie at the Victor Chang Cardiac Research Institute (Darlinghurst, New South Wales, Australia). We thank Dr. Lisa Conti at Quinnipiac University (Hamden, Connecticut, United States) for critical guidance on the PPI experiments, and members of the MacDonald lab for critical reading and feedback on the manuscript. Funding was provided by a Collaboration for Unprecedented Success and Excellence Grant (Syracuse University; to JLM) with additional laboratory support provided by the National Institutes of Health (JLM) [Grant number 1R01NS106285]. Declarations of interest: none

